# Dimension reduction of microbiome data linked *Bifidobacterium* to allergic rhinitis

**DOI:** 10.1101/2023.07.18.548236

**Authors:** Shohei Komaki, Yukari Sahoyama, Tsuyoshi Hachiya, Keita Koseki, Yusuke Ogata, Fumiaki Hamazato, Manabu Shiozawa, Tohru Nakagawa, Wataru Suda, Masahira Hattori, Eiryo Kawakami

## Abstract

Dimension reduction has been used to visualise the distribution of multidimensional microbiome data, but the composite variables calculated by the dimension reduction methods have not been widely used to investigate the relationship of the human gut microbiome with lifestyle and disease. In the present study, we applied several dimension reduction methods, including principal component analysis (PCA), principal coordinate analysis (PCoA), non-metric multidimensional scaling (NMDS), and non-negative matrix factorization (NMF), to a microbiome dataset from 186 subjects with symptoms of AR and 106 controls. All the dimension reduction methods supported that the enterotype clusters were overlapped in the dimension reduction plots, and that the distribution of microbial data points appeared to be continuous rather than discrete. Comparison of the composite variables calculated from the different dimension reduction methods showed that the characteristics of the composite variables differed between the dimension reduction methods. The second composite variable calculated from PCoA was significantly associated with the intake of several nutrients, including omega-3 polyunsaturated fatty acids, and the risk of AR. The composite variable was also correlated with the relative abundance of *Bifidobacterium*, and thus, *Bifidobacterium* was related to the risk of AR and intake of several nutrients through dimension reduction. Our results highlight the usefulness of the dimension reduction methods for investigating the association of microbial composition with lifestyle and disease in clinical research.

## Introduction

The human gut harbours a wide variety of microorganisms and the composition of the human microbiome varies from person to person. Growing evidence suggests that the microbial composition of the gut is associated with the lifestyle and health status of the host ^1^. In clinical research, enterotyping has been used as a method to characterise the microbial composition of the gut ^2,3^. In a seminal paper defining enterotypes for the first time, the human gut microbiota was classified into three enterotypes: P-type (*Prevotella*-rich), B-type (*Bacteroides*-rich), and R-type (*Ruminococcus*-rich) ^3^. Subsequent studies have shown that the composition of the human microbial community is not discretely distributed as enterotypes, but rather continuously distributed in typical populations ^4,5^.

Dimension reduction methods such as principal component analysis (PCA) and principal coordinate analysis (PCoA) are commonly used to visualise the distribution of the human microbiome community ^6,7^. Dimension reduction methods illustrate the distributions of microbial samples by mapping multidimensional data of microbial composition onto a few composite dimensions based on the distances (or dissimilarities) between samples. The composite variables calculated by the dimension reduction methods are robust to technical and biological noise. However, in clinical research, the composite variables calculated by dimension reduction methods have not been widely used to study the association of the human gut microbiome with lifestyle and disease.

There are several methods for calculating the dissimilarity between samples and obtaining composite variables from the distance matrix ^8^. Here, we investigated the association of the human gut microbial composition with dietary intake of 42 nutrients and the symptom of allergic rhinitis (AR), focusing on the application and comparison of several dimension reduction methods, including PCA, PCoA, non-metric multidimensional scaling (NMDS), and non-negative matrix factorization (NMF).

## Materials and methods

### Samples and datasets

In the present study, we re-analysed a microbiome dataset from 186 participants with symptoms of AR and 106 controls without symptoms of AR at the Hitachi Health Care Centre in Japan. The dataset was used in our previous study to identify up- and down-regulated microbial genera in AR patients compared to controls ^9^. No dimension reduction method was used in the previous study. The present study investigated the association of the human gut microbial composition with dietary intake of 42 nutrients and the symptom of AR, focusing on the application and comparison of dimension reduction methods.

In brief, food consumption data for the study participants were obtained using the Brief Self-Administered Diet History Questionnaire (BDHQ) and adjusted by energy using the density method ^10–12^. Bacterial DNA was isolated from faecal samples, followed by amplification of the 16S V1-V2 region by polymerase chain reaction (PCR) ^13^. An equal amount of each PCR amplicon was mixed and subjected to multiplex amplicon sequencing using MiSeq (2 × 300 paired-end). Filtered reads with BLAST match lengths < 90% to the representative sequence in the 16S databases, including the Ribosomal Database Project (RDP) (Release 11, Update 5), CORE (updated 13 October 2017; http://microbiome.osu.edu/), and a reference genome sequence database obtained from the NCBI FTP site (ftp://ftp.ncbi.nih.gov/genbank/, April 2013), were considered chimeras and removed. From the filtered reads, 3,000 high quality reads per sample were randomly selected. The total reads (number of samples × 3,000) were then sorted by the frequency of redundant sequences and grouped into operational taxonomic units (OTUs) using UCLUST with a sequence identity threshold of 97%. The representative sequences of the generated OTUs were subjected to a homology search against the above databases using the GLSEARCH program for taxonomic assignments. Phylum, genus and species level assignments were made using sequence similarity thresholds of 70%, 94% and 97%, respectively.

This study was approved by the Hitachi Hospital Group Ethics Committee (Approved No. 2018-5, 2019-10, and 2020-88), the Institutional Review Board of the Hitachi Ltd. (Approved No. 220-1 and 238-1), and the Research Ethics Committee (Approved No. H30-5). Written informed consent was obtained from all participants. This study was conducted in accordance with the principles of the Declaration of Helsinki.

### Dimension reduction of microbial composition

Genus-level abundance was expressed as a percentage. Genera with a mean relative abundance of ≥0.1% were included, resulting in a genus-level abundance matrix with 292 rows (samples) and 42 columns (genera).

We first performed enterotyping to classify the microbiome of 292 individuals as in the landmark study ^3^. Following the enterotyping R tutorial (https://enterotype.embl.de/), the Jensen-Shannon divergence (square root of the Jensen-Shannon distance) between all pairs of 292 samples was calculated and a 292 × 292 pairwise distance matrix was generated. The Jensen-Shannon distance is a symmetrized and smoothed version of the Kullback-Leibler divergence which measures the similarity of the probability distributions of two samples ^14^. Based on the distance matrix, 292 samples were then clustered into the discrete enterotypes by the partitioning around medoids algorithm using the clusterSim R package (version 0.50.1) ^15^. The number of clusters was set at 3 as in the seminal study ^3^.

We also used several dimension reduction methods, including PCA, PCoA, NMDS, and NMF, to obtain the continuous composite variables from the above-mentioned genus-level abundance matrix of 292 samples and 42 genera. The top 3 composite variables for each dimension reduction method were obtained for subsequent analyses.

To perform the PCA, we used the prcomp function implemented in the stats R package (version 4.2.1) ^16^ with the scale=TRUE option, which normalises the genus-level abundance matrix so that each column has a mean of 0 and a standard deviation of 1.

To perform the PCoA, we applied the cmdscale function in the stats R package (version 4.2.1) to the above-mentioned Jansen-Shannon distance to obtain the top 3 composite variables. In addition, we calculated the Bray-Curtis dissimilarity matrix ^17^ using the beta.pair.abund function implemented in the betapart R package, which measures the compositional difference between two ecological communities (version 1.5.6) ^18^ with “bray” specified as index.family. The cmdscale function was applied to the Bray-Curtis dissimilarity matrix to obtain the top 3 composite variables.

For the NMDS, the microbiome abundance matrix of 292 samples was standardised and multiplied by the total sample size using the decostand function from the vegan R package (version 2.6.2) ^19^. The NMDS analysis was then performed to obtain the top 3 composite variables using the metaMDS function from the vegan R package with the distance index “bray”, the number of dimensions (k) 3, the maximum number of random starts in search of a stable solution (maxit) 100.

Non-negative matrix factorization is a dimension reduction method that decomposes a non-negative matrix V into two non-negative matrices W and H, such that V is approximately equal to W multiplied by H ^20^. The matrix W represents the composite variables, while H represents the coefficients when the original data are expressed as a linear combination of the composite variables. The number of composite variables is set as the rank of W and H. The reconstruction error between V and WH is minimised by iteratively updating W and H according to some loss function. We used the NNLM R package (version 0.4.4) ^21^ to apply NMF to the genus-level abundance matrix V. We used the Kullback-Leibler divergence as the loss function to minimise the reconstruction error. To determine the optimal rank, we used the masking approach through missing value imputation proposed in previous studies ^22,23^. Some elements in the original matrix were randomly replaced by NA and masked to obtain the noisy matrix. The reconstruction error is calculated only for the replaced elements. In this study, we calculated the median reconstruction error, obtained from 50 independent decompositions for ranks from 2 to 10. The reconstruction error was minimised when the rank was set to 3. For subsequent analysis, we used W at rank 3 representing composite variables.

For comparison, we also calculated the ratio of *Prevotella* to *Bacteroides* (P/B ratio) ^24,25^.

### Statistical analysis

To examine the similarity between composite variables, and to examine the association between composite variables and genus-level abundance, we applied the pairwise Spearman’s rank correlation test using the cor function from the stats R package with the “spearman” method. We evaluated the association of composite variables with the intake levels of 42 nutrients using the Spearman’s rank correlation test. The association of composite variables with AR was tested by the Wilcoxon-Mann-Whitney test using the wilcox_test function from the coin R package (version 1.4.2) ^26^, with the distribution set to “exact”. The resulting *P*-values were corrected for multiple testing in each dimension reduction method by the Benjamini and Hochberg false discovery rate (FDR) ^27^ using the p.adjust function in the stats R package, specifying “BH” as method.

## Results

### Continuous distribution of gut microbial community

There were 42 genera with mean relative abundance ≥0.1%. The Jensen-Shannon divergence matrix was calculated from the 42-dimensional genus-level data of the 292 individuals. Enterotypes were calculated from the Jensen-Shannon divergence matrix (**Figure 1a**). *Bacteroides* and *Prevotella* were abundant in the enterotype 1 (corresponding to B-type) and 3 (P-type), respectively (**Figure 1b; Table 1**). The abundance of *Ruminococcus* was similar between enterotypes, whereas *Bifidobacterium* was abundant in the enterotype 2 (**Figure 1b; Table 1**). The distributions of enterotypes 1 and 2 overlapped in the PCoA plots, emphasising that the microbial composition was continuous rather than discrete. The continuous distribution of gut microbial composition was further supported by other dimension reduction methods, including PCA (**Figure 2a**), PCoA with the Bray-Curtis dissimilarity (**Figure 2b**), NMDS (**Figure 2c**), and NMF (**Figure 2d**).

**Table 1.**
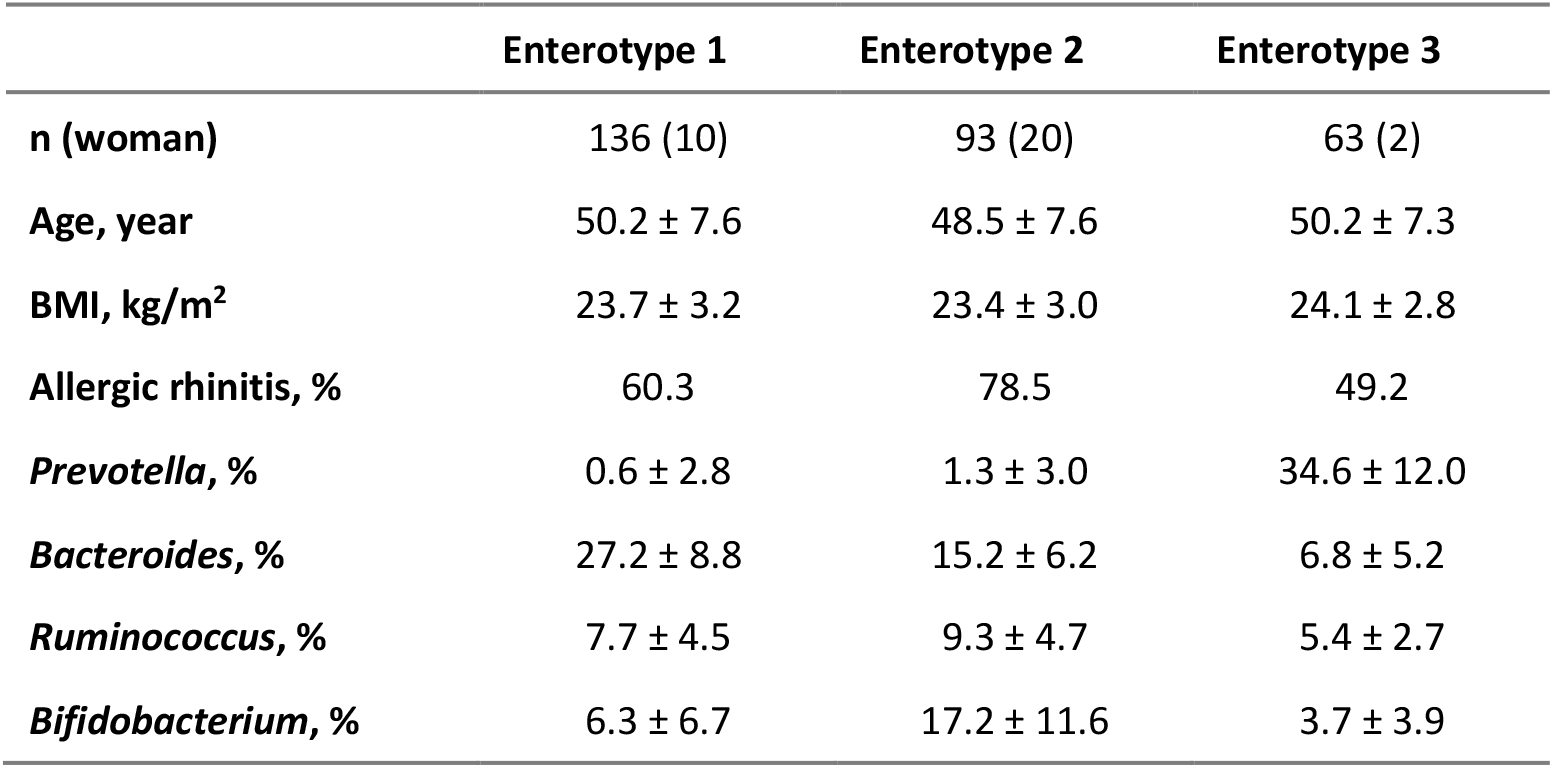
Characteristics of the study participants.

**Figure 1.**
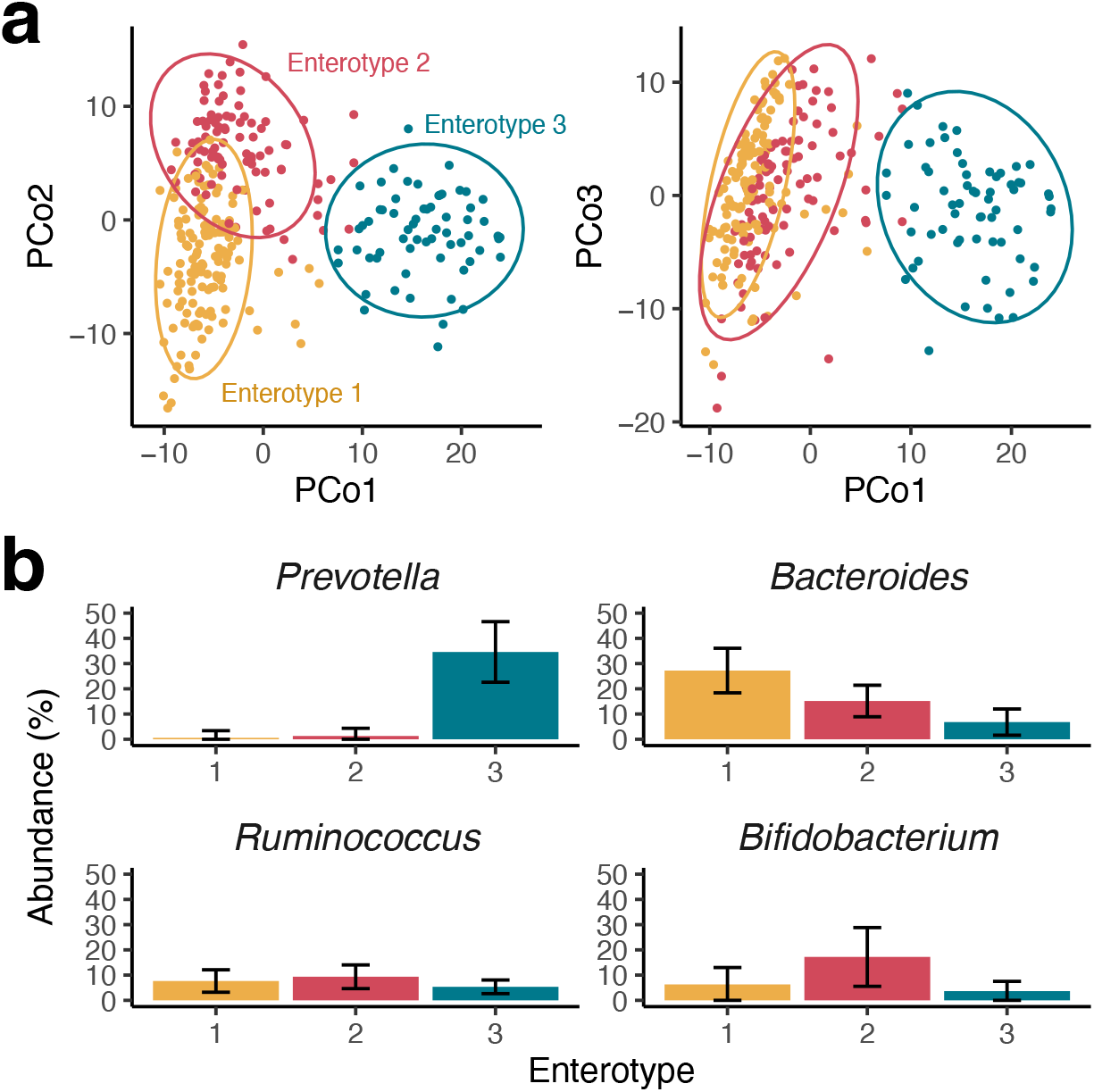
Enterotypes and distribution of gut microbial composition. (a) Principal coordinate analysis plots based on the Jensen-Shannon divergence. The left panel shows the top 1 (x-axis) and 2 (y-axis) composite variables, while the right panel shows the top 1 (x-axis) and 3 (y-axis) composite variables. The dot colour indicates the enterotype calculated from the same Jensen-Shannon divergence matrix. (b) Relative abundances of representative genera by enterotype.

**Figure 2.**
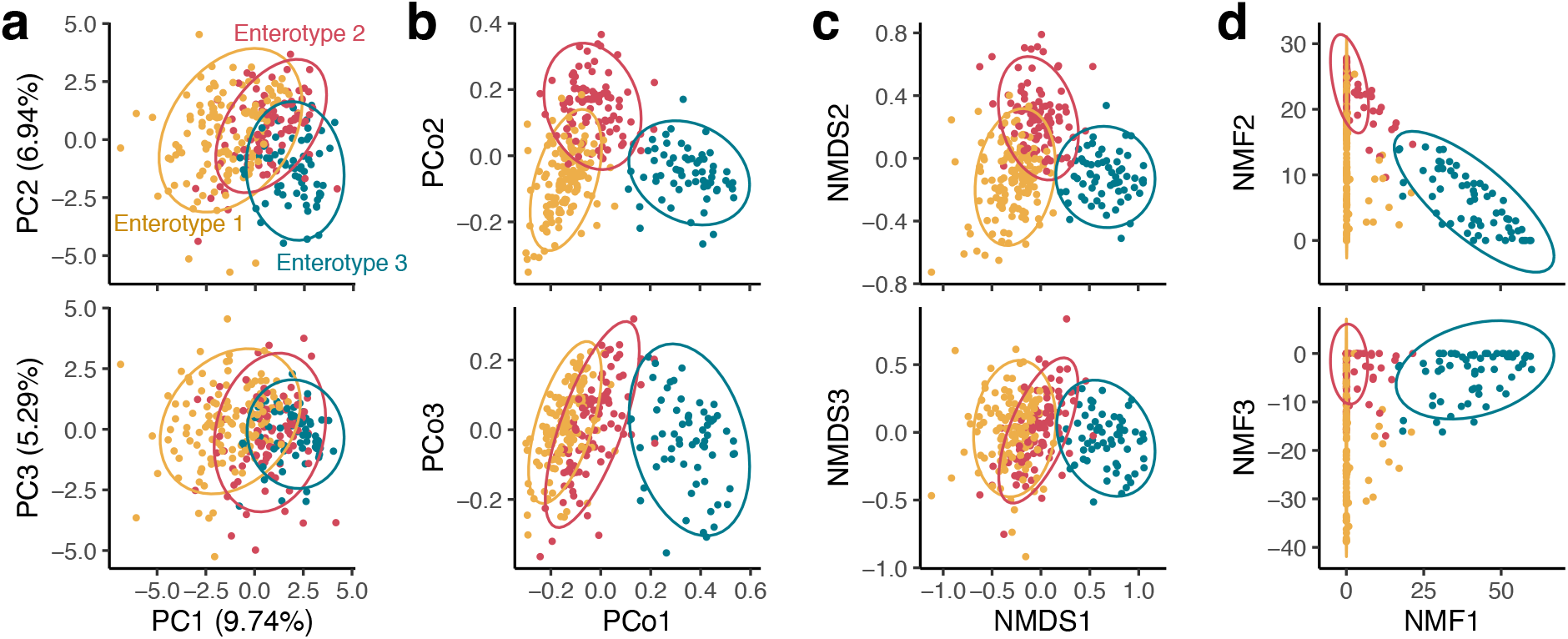
Distribution of the gut microbial community using different dimension reduction methods. (a) principal component analysis, (b) principal coordinate analysis with the Bray-Curtis distance, (c) non-metric multidimensional scaling with the Bray-Curtis distance, and (d) non-negative matrix factorization. The top panel shows the top 1 (x-axis) and 2 (y-axis) composite variables, while the bottom panel shows the top 1 (x-axis) and 3 (y-axis) composite variables. The colour of the point indicates the enterotype.

### Comparison of composite variables calculated using different dimension reduction methods

To compare the composite variables obtained from the different dimension reduction methods, we calculated the Spearman’s rank correlation between pairs of composite variables (**Figure 3a**). The results showed that the top 3 composite variables calculated from the PCoA using the Jensen-Shannon distance were highly correlated with those calculated from the PCoA and NMDS using the Bray-Curtis distance. The top 1 composite variable calculated from PCA was highly correlated with those from PCoA and NMDS, while the second and third composite variables from PCA were not remarkably correlated with those calculated from the other methods. The first composite variable calculated by NMF was highly correlated with the P/B ratio, and the second and third composite variables calculated by NMF were highly correlated with each other.

**Figure 3.**
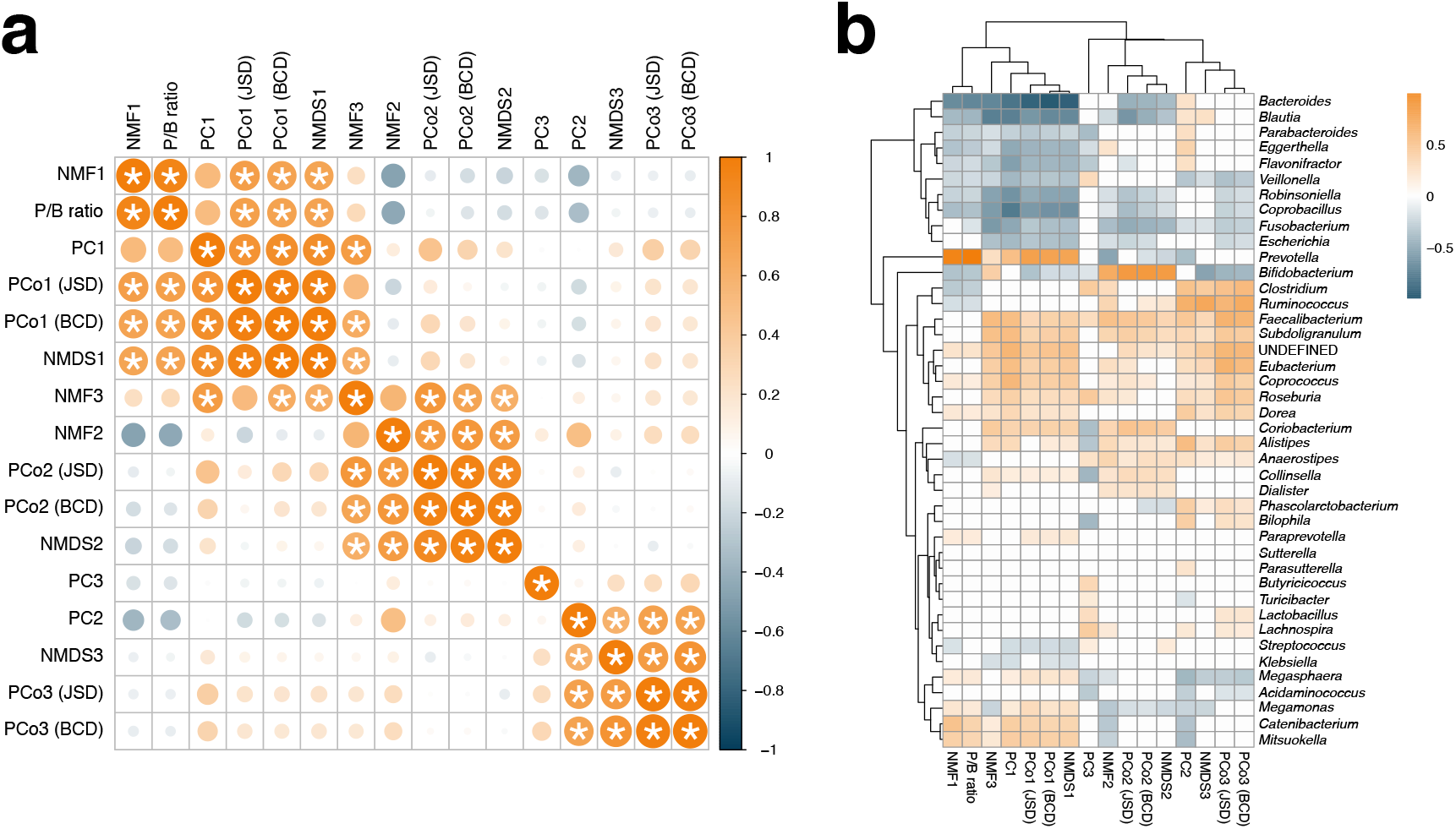
Comparison and characterisation of composite variables. (a) Pairwise Spearman’s rank correlations between composite variables. X- and y-axes were ordered based on hierarchical clustering. Highly correlated pairs (Spearman’s rank correlation >0.8) are indicated by asterisks (*). (b) Spearman’s rank correlations between composite variable and genus-level relative abundance. Non-significant pairs (FDR-adjusted *P*-value >0.05) are shown as white cells. NMF: non-negative matrix factorization, PB ratio: *Prevotella*-*Bacteroides* ratio, PC: principal component, PCo: principal coordinate, NMDS: non-metric multidimensional scaling, JSD: Jensen-Shannon divergence, BCD: Bray-Curtis distance.

The first composite variable calculated using PCoA and NMDS was positively correlated with the relative abundance of *Prevotella* and negatively correlated with *Bacteroides* (**Figure 3b**). The second composite variable calculated using PCoA and NMDS was positively correlated with *Bifidobacterium*, while the third composite variable was negatively correlated with *Bifidobacterium*. The first composite variable calculated using NMF was positively correlated with *Prevotella* and negatively correlated with *Bacteroides*, as was the P/B ratio.

### Nutrients and AR were related to microbial composition via composite variables

We examined the association of composite variables calculated using different dimension reduction methods with the consumption levels of 42 nutrients and the risk of AR (**Figure 4**). The risk of AR was associated with the second composite variable calculated from the Bray-Curtis distance (PCoA and NMDS) (subjects with symptoms had larger composite variables; FDR adjusted *P*-value < 0.05), while the other composite variables were not significantly associated with the risk of AR. The second composite variable calculated from the Bray-Curtis distance (PCoA), which was positively correlated with the relative abundance of *Bifidobacterium*, was also associated with the intake of several nutrients, including carbohydrate [CHO], saturated fat [SFA], n-3 polyunsaturated fat [N3], eicosapentaenoic acid [C205n3], vitamin B12 [VB12], and alcohol [ALC] (**Figure 4b**). Carbohydrate [CHO], saturated fat [SFA], and alcohol [ALC] were also associated with the second composite variables calculated using the Jensen-Shannon divergence (**Figure 4a**), NMDS (**Figure 4c**), and NMF (**Figure 4e**), whereas no nutrient was associated with the composite variables calculated using PCA (FDR adjusted *P*-value > 0.05) (**Figure 4d**), nor was P/B ratio (**Figure 4f**).

**Figure 4.**
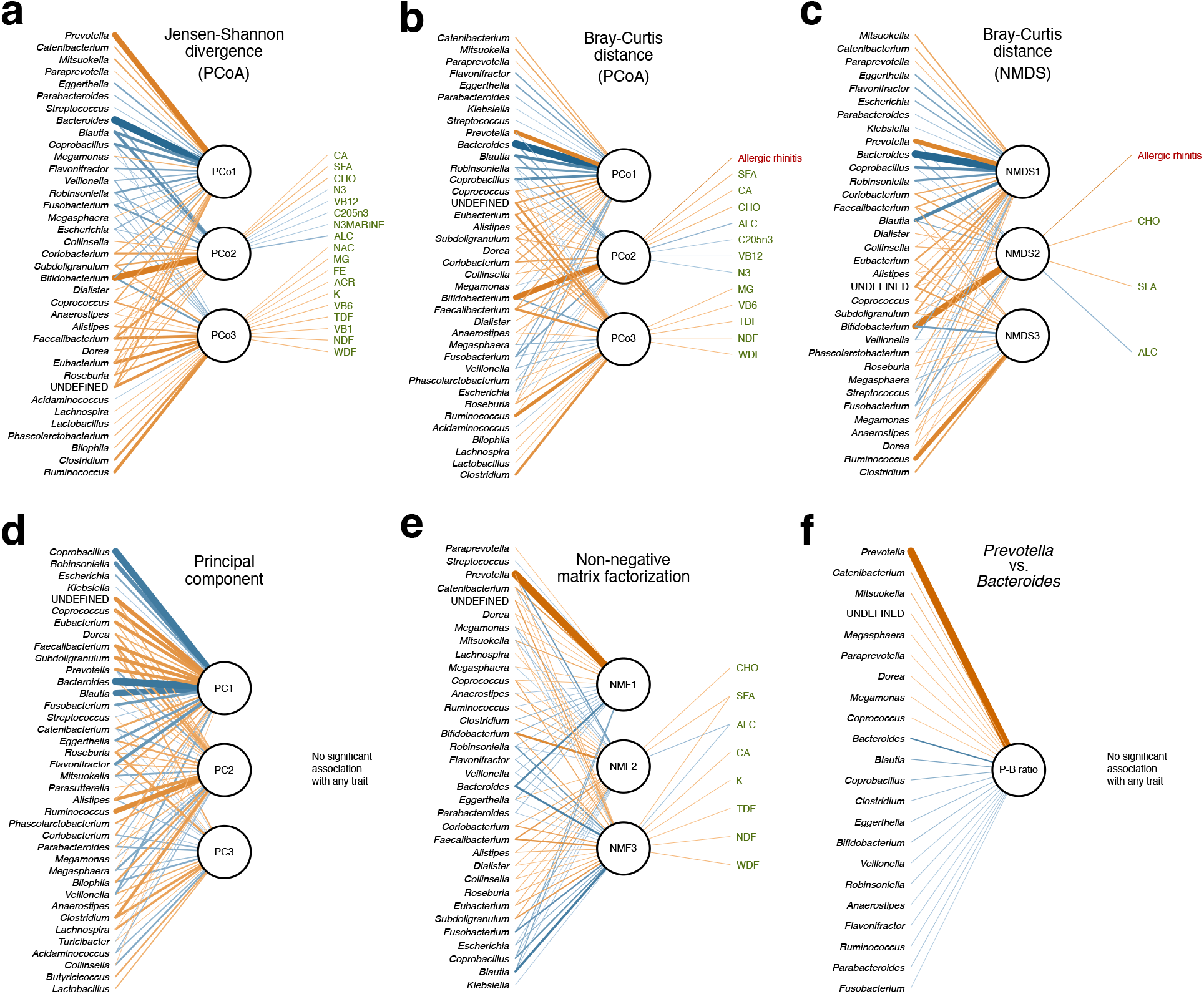
Associations of microbial composition with nutrient intakes and AR via composite variables. Circles indicate the top 3 composite variables calculated using different dimension reduction methods. Significant associations (FDR adjusted *P*-value < 0.05) of the composite variables with genus-level abundance, nutrient intake, and the symptom of AR are shown. Line thickness indicates the strength of association. Orange and blue line indicate positive and negative associations, respectively. (a) Principal coordinate analysis with the Jensen-Shannon divergence, (b) principal coordinate analysis with the Bray-Curtis distance, (c) non-metric multidimensional scaling with the Bray-Curtis distance, (d) principal component analysis, (e) non-negative matrix factorization, and (f) *Prevotella* to *Bacteroides* ratio. ACR: α-carotene, ALC: alcohol, C205n3: eicosapentaenoic acid, CA: calcium, CHO: carbohydrate, FE: iron, K: potassium, MG: magnesium, N3: n-3 polyunsaturated fat, N3MARINE: n-3 polyunsaturated fat of marine origin a, NAC: niacin, NDF: insoluble dietary fibre, SFA: saturated fat, TDF: total dietary fibre, VB1: thiamin, VB12: vitamin B12, VB6: vitamin B6, WDF: soluble dietary fibre.

The direct comparison between genus-level relative abundance and nutrient consumption level (43 genera × 42 nutrients) showed only one significant association after correction for multiple testing (*Fusobacterium* and alcohol, adjusted *P*=0.02). In this pairwise comparison, the association between *Bifidobacterium* and nutrients was not significant.

## Discussion

Dimension reduction has been used to visualise the distribution of multidimensional microbiome data ^6,7^, but the composite variables calculated by the dimension reduction methods have not been widely used to investigate the relationship of the human gut microbiome with lifestyle and disease. In the present study, we applied several dimension reduction methods, including PCA, PCoA, NMDS, and NMF, to a microbiome dataset from 186 subjects with symptoms of AR and 106 controls. All the dimension reduction methods supported that the enterotype clusters were overlapped in the dimension reduction plots, and that microbial composition appeared to be continuous rather than discrete. The top 3 composite variables calculated from PCoA and NMDS were highly correlated, whereas the top 3 composite variables from other methods were not corresponded, suggesting that the characteristics of the composite variables differed between the dimension reduction methods.

The second composite variables calculated from PCoA and NMDS were significantly associated with the *Bifidobacterium* abundance and also with the several nutrient intakes and the risk of AR, linking *Bifidobacterium* with nutrient intake and risk of AR, whereas the direct comparison found no significant association between them. The result highlights that dimension reduction methods are useful for finding novel associations by reducing the number of statistical tests and relaxing the *P*-value threshold after correction for multiple testing.

The genus *Bifidobacterium* has been recognized to confer health benefits to the host also through its interaction with the host’s immune system ^28,29^. These benefits comprise both local effects, which result from the contribution of *Bifidobacterium* to the intestinal barrier function—which ultimately translates into systemic health—and systemic effects, which stem from the microorganism’s impact on specific pathways through extracellular structures and metabolites. An example of this is the anti-inflammatory response that is elicited by acetate produced by *Bifidobacterium* ^30^. *Bifidobacterium* can digest complex carbohydrates, such as glucans, into acetate which is further digested into butyrate by other gut microorganisms. Butyrate is known to possess anti-inflammatory properties that include the production of TGF-β, IL-18, and IL-10 cytokines by antigen-presenting cells and IECs, which together stimulate the differentiation of naïve T cells into Treg cells. Additionally, *Bifidobacterium* produce two types of pili, hair-like structures found on the surface of bacteria, and pili produced in certain bifidobacterial strains have been shown to stimulate TNF-α levels in macrophages while suppressing other pro-inflammatory cytokines that are associated with systemic immune responses ^31^.

The association between *Bifidobacterium* and allergic symptoms has indeed been reported. For example, an observational microbiome study reported that patients with atopy and asthma tended to have a lower abundance of *Bifidobacterium* ^32^. Although studies on the relationship between allergic rhinitis and the intestinal *Bifidobacterium* abundance are limited, one study reported that the symptom of allergic rhinitis was reduced by oral administration of probiotic *B. lactis* ^33^. However, the increase in intestinal *Bifidobacterium* was not confirmed. In addition, it is debated whether *Bifidobacterium* is beneficial or not, and at least, the genus *Bifidobacterium* is not uniformly beneficial ^34^.

There are several limitations to this study. First, we did not evaluate the performance of the different dimension reduction methods. We applied several dimension reduction methods to a case-control dataset and examined the association of the composite variables calculated using different dimension reduction methods with nutrient intake and the risk of AR. Further studies are needed to investigate whether dimension reduction methods can contribute to an increased ability to distinguish patients from controls on the basis of discriminatory power. Second, we only analysed one microbiome dataset. Further research using a variety of microbiome datasets is warranted to determine the advantages and disadvantages of dimension reduction methods. For example, our data showed that the first composite variable calculated by NMF was positively correlated with the P/B ratio, suggesting that NMF may be able to extract a meaningful dimension from microbiome data alone without prior knowledge.

In conclusion, *Bifidobacterium* was associated with intake of multiple nutrients and the risk of AR by applying multiple dimension reduction methods to a microbiome dataset. Our results highlight the usefulness of the dimension reduction methods for investigating the association of microbial composition with lifestyle and disease in clinical research.

## Data availability

The data are not available for public access because of participant privacy concerns, but are available from the corresponding author on reasonable request.

## Acknowledgements

This research did not receive any specific grant from funding agencies in either the public, commercial, or not-for-profit sectors.

## Author contributions

S.K., T.H., K.K., and E.K. wrote the manuscript. T.N. oversaw clinical data management. Y.S., T.H., F.H., and M.S. managed the nutritional data. Y.O., W.S., and M.H. managed the microbiome data. S.K., Y.S., T.H., K.K., and E.K. performed the statistical analysis. F.H., M.S., M.H., and E.K. supervised the work. Y.S., F.H., M.S., T.N., M.H., and E.K. designed and coordinated the project. All authors commented on and approved the manuscript.

## References

1. Hou, K. et al. Microbiota in health and diseases. Signal Transduction and Targeted Therapy vol. 7 Preprint at https://doi.org/10.1038/s41392-022-00974-4 (2022).

2. Wu, G. D. et al. Linking Long-Term Dietary Patterns with Gut Microbial Enterotypes. Science (1979) 334, 105–108 (2011).

3. Arumugam, M. et al. Enterotypes of the human gut microbiome. Nature 473, 174–180 (2011).

4. Mobeen, F., Sharma, V. & Prakash, T. Enterotype Variations of the Healthy Human Gut Microbiome in Different Geographical Regions. Bioinformation 14, 560–573 (2018).

5. Knights, D. et al. Rethinking enterotypes. Cell Host and Microbe vol. 16 433–437 Preprint at https://doi.org/10.1016/j.chom.2014.09.013 (2014).

6. Yang, S. et al. The gut microbiome and antibiotic resistome of chronic diarrhea rhesus macaques (Macaca mulatta) and its similarity to the human gut microbiome. Microbiome 10, (2022).

7. Zhu, L. et al. Gut microbial characteristics of adult patients with allergy rhinitis. Microb Cell Fact 19, (2020).

8. Cheng, M. & Ning, K. Stereotypes About Enterotype: the Old and New Ideas. Genomics, Proteomics and Bioinformatics vol. 17 4–12 Preprint at https://doi.org/10.1016/j.gpb.2018.02.004 (2019).

9. Sahoyama, Y. et al. Multiple nutritional and gut microbial factors associated with allergic rhinitis: the Hitachi Health Study. Sci Rep 12, (2022).

10. Kobayashi, S. et al. Comparison of relative validity of food group intakes estimated by comprehensive and brief-type self-administered diet history questionnaires against 16 d dietary records in Japanese adults. Public Health Nutr 14, 1200–1211 (2011).

11. Kobayashi, S. et al. Both comprehensive and brief self-administered diet history questionnaires satisfactorily rank nutrient intakes in Japanese adults. J Epidemiol 22, 151–159 (2012).

12. Willett, W. & Stampfer, M. J. Total energy intake: implications for epidemiologic analyses. Am J Epidemiol 124, 17–27 (1986).

13. Kim, S. W. et al. Robustness of gut microbiota of healthy adults in response to probiotic intervention revealed by high-throughput pyrosequencing. DNA Research 20, 241–253 (2013).

14. Endres, D. M. & Schindelin, J. E. A new metric for probability distributions. IEEE Transactions on Information Theory vol. 49 1858–1860 Preprint at https://doi.org/10.1109/TIT.2003.813506 (2003).

15. Walesiak, M. & Dudek, A. The Choice of Variable Normalization Method in Cluster Analysis. in Education Excellence and Innovation Management: A 2025 Vision to Sustain Economic Development During Global Challenges (ed. Soliman, K. S.) 325–340 (International Business Information Management Association (IBIMA), 2020).

16. R Core Team. R: A language and environment for statistical computing. Preprint at (2022).

17. Baselga, A. Separating the two components of abundance-based dissimilarity: Balanced changes in abundance vs. abundance gradients. Methods Ecol Evol 4, 552–557 (2013).

18. Baselga, A. et al. betapart: Partitioning Beta Diversity into Turnover and Nestedness Components. Preprint at (2022).

19. Oksanen, J. et al. vegan: Community Ecology Package. Preprint at https://CRAN.R-project.org/package=vegan (2022).

20. Lee, D. D. & Seung, H. S. Learning the parts of objects by non-negative matrix factorization. Nature 401, 788–791 (1999).

21. Lin, X. & Paul C Boutros. NNLM: Fast and Versatile Non-Negative Matrix Factorization. Preprint at https://github.com/linxihui/NNLM (2020).

22. Ikeda, K. et al. Detecting time-evolving phenotypic components of adverse reactions against BNT162b2 SARS-CoV-2 vaccine via non-negative tensor factorization. iScience 25, 105237 (2022).

23. Lin, X. & Boutros, P. C. Optimization and expansion of non-negative matrix factorization. BMC Bioinformatics 21, (2020).

24. Levy, R. et al. Longitudinal analysis reveals transition barriers between dominant ecological states in the gut microbiome. Proceedings of the National Academy of Sciences 117, 13839–13845 (2020).

25. Hjorth, M. F. et al. Prevotella-to-Bacteroides ratio predicts body weight and fat loss success on 24-week diets varying in macronutrient composition and dietary fiber: results from a post-hoc analysis. Int J Obes 43, 149–157 (2019).

26. Hothorn, T., Hornik, K., van de Wiel, M. A. & Zeileis, A. A Lego System for Conditional Inference. Am Stat 60, 257–263 (2006).

27. Benjamini, Y. & Hochberg, Y. Controlling the False Discovery Rate: A Practical and Powerful Approach to Multiple Testing. Journal of the Royal Statistical Society. Series B (Methodological) 57, 289–300 (1995).

28. Ruiz, L., Delgado, S., Ruas-Madiedo, P., Sánchez, B. & Margolles, A. Bifidobacteria and their molecular communication with the immune system. Frontiers in Microbiology vol. 8 Preprint at https://doi.org/10.3389/fmicb.2017.02345 (2017).

29. Alessandri, G., Ossiprandi, M. C., MacSharry, J., van Sinderen, D. & Ventura, M. Bifidobacterial Dialogue With Its Human Host and Consequent Modulation of the Immune System. Frontiers in Immunology vol. 10 Preprint at https://doi.org/10.3389/fimmu.2019.02348 (2019).

30. Fukuda, S. et al. Bifidobacteria can protect from enteropathogenic infection through production of acetate. Nature 469, 543–549 (2011).

31. Duranti, S. et al. Elucidating the gut microbiome of ulcerative colitis: bifidobacteria as novel microbial biomarkers. FEMS Microbiol Ecol 92, fiw191 (2016).

32. Fujimura, K. E. et al. Neonatal gut microbiota associates with childhood multisensitized atopy and T cell differentiation. Nat Med 22, 1187–1191 (2016).

33. Singh, A. et al. Immune-modulatory effect of probiotic Bifidobacterium lactis NCC2818 in individuals suffering from seasonal allergic rhinitis to grass pollen: An exploratory, randomized, placebo-controlled clinical trial. Eur J Clin Nutr 67, 161–167 (2013).

34. Hidalgo-Cantabrana, C. et al. Bifidobacteria and Their Health-Promoting Effects. Microbiol Spectr 5, (2017).

